# Multimodal epi-mode 3D quantitative phase and fluorescence microscopy enables supervised volumetric virtual fluorescence in bulk tissue

**DOI:** 10.64898/2026.10.21.760751

**Authors:** Zhenmin Li, Srinidhi Bharadwaj, Aaron D Silva Trenkle, Daniel Houston, Kyan Nguyen, Nimai Ponna, Jacob P. Callaway, Gabriel A. Kwong, Francisco E. Robles

## Abstract

We introduce a multimodal microscope that combines quantitative oblique back-illumination microscopy (qOBM) with wide-field epi-fluorescence deconvolution microscopy to acquire intrinsically co-registered phase and fluorescence volumes of fresh, bulk tissue. Using paired qOBM phase and DAPI fluorescence images of fresh *ex vivo* mouse brain tissue, we train a supervised conditional generative adversarial network (cGAN) to predict DAPI-like nuclear contrast from the structural information encoded in the phase images. Results not only demonstrate high-fidelity image translation but also show improved imaging depth and more homogeneous staining. Additionally, we combine the qOBM phase contrast with captured or virtual fluorescence nuclear images to produce volumetric pseudo-H&E representations of fresh tissue, enabling a supervised approach for achieving robust virtual histological staining. This work paves the way for virtual fluorescence imaging of many more structural targets and supervised virtual histological staining of thick, unaltered, bulk tissues *ex vivo* and potentially *in vivo*. Such capabilities could have significant implications for biological research as well as for point-of-care, intraoperative, and other clinical applications.

## 1 Introduction

Quantitative phase imaging (QPI) and fluorescence microscopy are complementary imaging technologies that enable, respectively, highly sensitive endogenous refractive index contrast that yields rich biophysical information [1], and high-contrast molecular imaging of labeled structures [2]. Each also has unique limitations: phase contrast lacks molecular specificity and cannot highlight specific structures of interest, whereas fluorescence microscopy suffers from photobleaching and phototoxicity and only provides contrast of targeted molecules or structures. To explore the unique insight provided by both label-free phase contrast and fluorescence imaging, researchers have developed multimodal approaches that combine these complementary techniques [3, 4]. More recently, these multimodal systems have been leveraged in efforts to replace the fluorescence channel(s) by learning a mapping from phase images to their labeled counterparts, thus providing the strengths of the two complementary technologies without their weaknesses [5–7]. While extremely promising, state-of-the-art methods that enable high-resolution virtual fluorescence staining of unlabeled specimens face significant limitations given traditional QPI methods are only able to image thin samples, such as cultured cells or sectioned tissues. While 3D fluorescence imaging of bulk tissues is readily achievable [8, 9], label-free virtual staining of such samples has remained elusive. This limitation significantly hampers the huge biological and clinical impact that virtual fluorescence staining could achieve.

In this work, we develop a multimodal microscope that combines quantitative oblique back-illumination microscopy (qOBM) with wide-field deconvolution epifluorescence microscopy to enable (1) co-registered QPI and fluorescence volumes of stained fresh, bulk tissue, and (2) phase-derived virtual staining of unstained tissues. qOBM is a recently developed label-free imaging platform that provides high-resolution, 3D quantitative phase images and 3D refractive index (RI) tomography of thick, scattering biological samples in epi-mode [10, 11], and because it uses epiillumination, it can be integrated seamlessly with epi-fluorescence. To date, qOBM has been applied to image a wide range of biological specimens *in vivo, ex vivo*, and *in vitro*, including skin [12–14], organoids [15, 16], brain [17, 18] and many more [19, 20], with imaging depth well beyond 160 *µ* m [21]. The multimodal system developed here is designed to be simple and cost-effective, requiring only minor modifications to a standard wide-field microscope. The fluorescence and phase images are inherently co-registered, ensuring precise 3D spatial alignment, which allows us to train a supervised conditional generative adversarial network (cGAN) to virtually stain label-free qOBM phase images with robust performance. Furthermore, we achieve pseudo and virtual hematoxylin and eosin (H&E) histology image representations by assigning the measured or virtual nuclear fluorescence as the nuclear stain (hematoxylin-like), and the phase image as the counter-stain (eosin-like). This work represents a significant step towards high-resolution, label-free 3D virtual fluorescence imaging and virtual H&E of bulk tissues with robust nuclear segmentation *ex vivo*. The approach could potentially be applied *in vivo*, which could have a broad range of research and clinical applications.

## 2 Results

### 2.1 Multimodal qOBM & fluorescence system design and performance

Our qOBM system integrates a conventional bright-field microscope with four multimode optical fibers that are evenly separated around the objective and are affixed at a 55-degree angle relative to the optical axis, optimized for oblique back-illumination [22]. The four fibers are connected to four 720 nm LEDs, which sequentially illuminate the sample (see Fig. 1**d**, where only one LED is turned on). The system uses a Nikon S Plan Fluor LWD 60× (0.7 NA) objective, a two-inch tube lens with 180 mm focal length (AC508-180-A, Thorlabs) and an sCMOS camera (pco.edge 4.2 LT). With 720 nm illumination and a 0.7 NA objective, the theoretical lateral and axial resolutions of qOBM are 627 nm and 2.94 *µ*m, respectively.

**Fig. 1.**
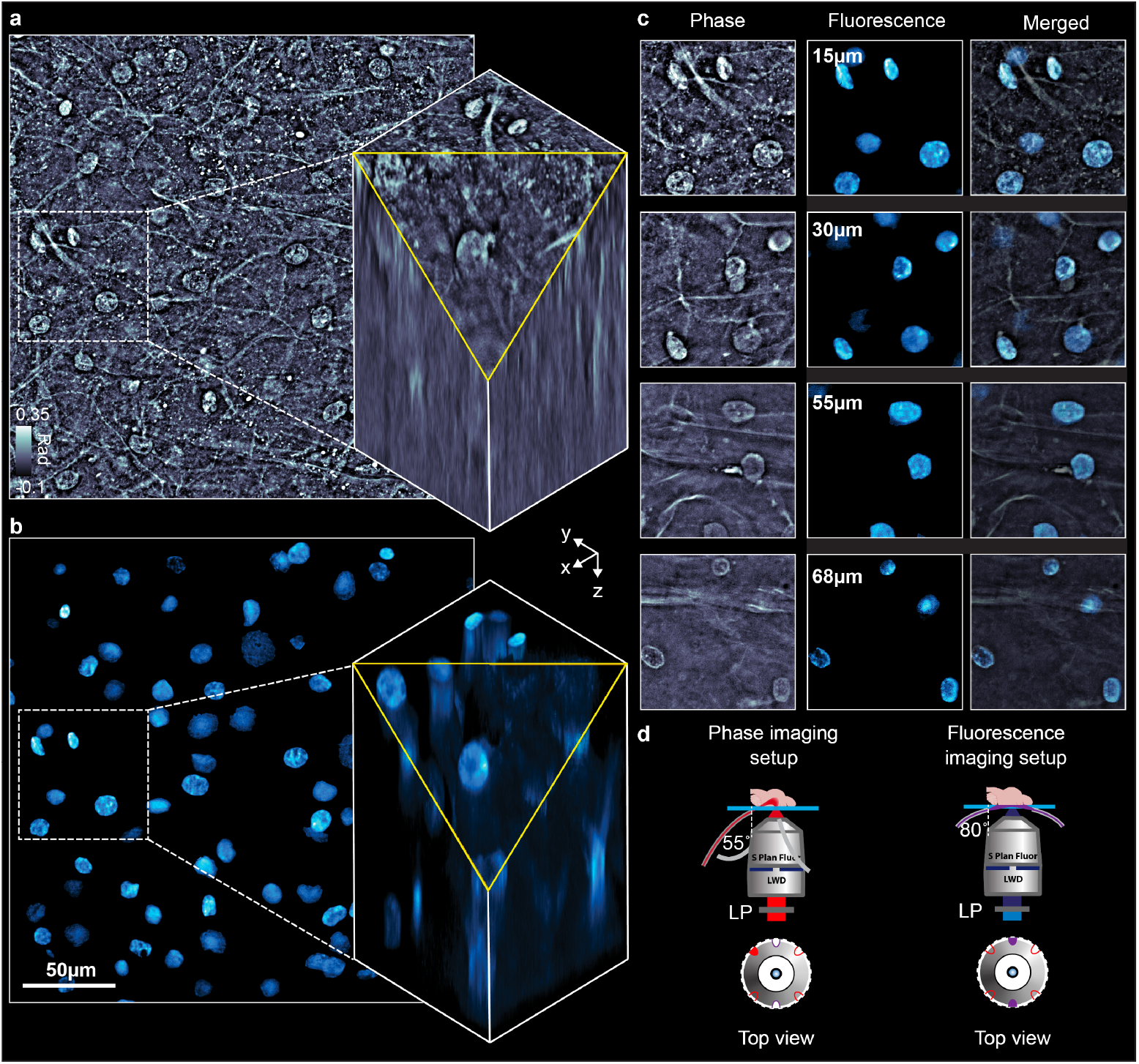
Multimodal imaging system configuration and capabilities. **(a)**: qOBM phase image of cervix and 3D phase volume of the square area indicated by the dashed line. **(b)**: DAPI fluorescence image and volume of the same FoV as in **a. (c)**: Depth-resolved cross-sections from the same ROI in **a** and **b**, demonstrating pixel-level registration of the two modalities across three dimensions. **(d)**: Multimodal system (side view and top view in top and bottom, respectively) during phase imaging (left) and fluorescence imaging mode (right).

To integrate the fluorescence imaging capabilities, we add two more multimode fibers connected to LEDs that have the desired excitation wavelength, in this case 365 nm to excite DAPI. The two fibers are positioned opposite each other around the objective and use a different illumination angle of 80-degree such that they illuminate the center of the tissue surface directly, instead of forming a banana curve to generate oblique back-illumination as with qOBM. For fluorescence imaging, the two LEDs turn on simultaneously to maximize the excitation light and provide even illumination across the field-of-view (see Fig. 1**d**). A long-pass filter (FELH0450, Thorlabs) with a cutoff wavelength of 450 nm is added on the detection path to remove the excitation light and allow detection of the fluorescence emission as well as the light used for qOBM. With DAPI emission peak at ∼461 nm, the corresponding theoretical lateral and axial resolutions of the fluorescence channel are 402 nm and 1.88 *µ*m, respectively. These modifications enable acquisition of fluorescence images with seamless integration with the existing qOBM phase imaging system. Using the same detection optics, five acquisitions are taken (four for qOBM and one for fluorescence) to acquire intrinsically co-registered quantitative phase and fluorescence images. Additional fluorescence channels can be included by simply adding more excitation wavelengths and/or rejection filters.

The system design is illustrated in Fig. 1**d**. Notably, the hardware modifications to a typical bright-field system are minimal, providing a substantially more accessible and versatile platform for combined volumetric phase/RI and fluorescence imaging than a recently reported reflection-mode approach that reconstructs RI from two-photon-excited fluorescence using a femtosecond laser and computationally intensive iterative neural-field reconstruction [23]. Moreover, because qOBM acquires phase independently of fluorescence, the platform also supports unlabeled specimens and enables phase-only virtual fluorescence inference after supervised training.

To validate the newly designed system, we used DAPI to visualize nuclei as a complementary imaging channel to phase imaging. DAPI binds specifically to DNA, enabling precise identification and localization of nuclei while also minimally altering the refractive index composition of tissue (see Supplemental Information, section 2). De-identified and discarded cervical tissue samples were provided by Emory University Hospital and imaged at the Georgia Institute of Technology. The tissues were stained for 1.5 hours and washed with phosphate-buffered saline (see Methods section for detailed protocol), then imaged with both modalities. For each field-of-view (FoV), we performed a z-stack, where at each depth we first acquired a qOBM image (with a typical exposure of 50 ms per one of four captures) and then the fluorescence image (typical exposure of 450 ms) before moving to the next depth. The net frame rate for the qOBM phase + fluorescence multimodal images was ∼1 Hz. The size of the FoV with the 60× is 243 × 243 *µ*m^2^. Acquiring z-stack allowed us to perform a 3D deconvolution (see Methods section) to improve the 3D fluorescence imaging results.

Figures 1**a** and 1**b** show a representative result. Here, the nuclei are visible in both modalities, and are well aligned across the volume in all dimensions. We note that the fluorescence images indeed have higher contrast for the nuclei–a strength of fluorescence microscopy–but the phase images still enable clear visualization of these structures. Figure 1**c** shows phase images and fluorescence images at four different depths along the stratified squamous epithelium. As we image deeper, nuclei become less dense, which is the natural maturation gradient of stratified squamous epithelium where cells mature and expand. Moreover, the examples in Fig. 1**c** and Fig. 2 show that the phase and fluorescence images exhibit similar sub-nuclear structures. This correspondence arises because DAPI binds specifically to DNA, while condensed, DNA-rich chromatin exhibits a modestly higher mass density, and thus higher refractive index, than the surrounding euchromatin [24]. Both modalities are therefore sensitive to the same underlying chromatin organization. Also, both imaging modalities experience depth-dependent signal-to-noise ratio (SNR) degradation, with the fluorescence channel showing a more rapid signal attenuation due to the shorter wavelengths of both excitation and emission light, as well as the limited penetration of the stain.

**Fig. 2.**
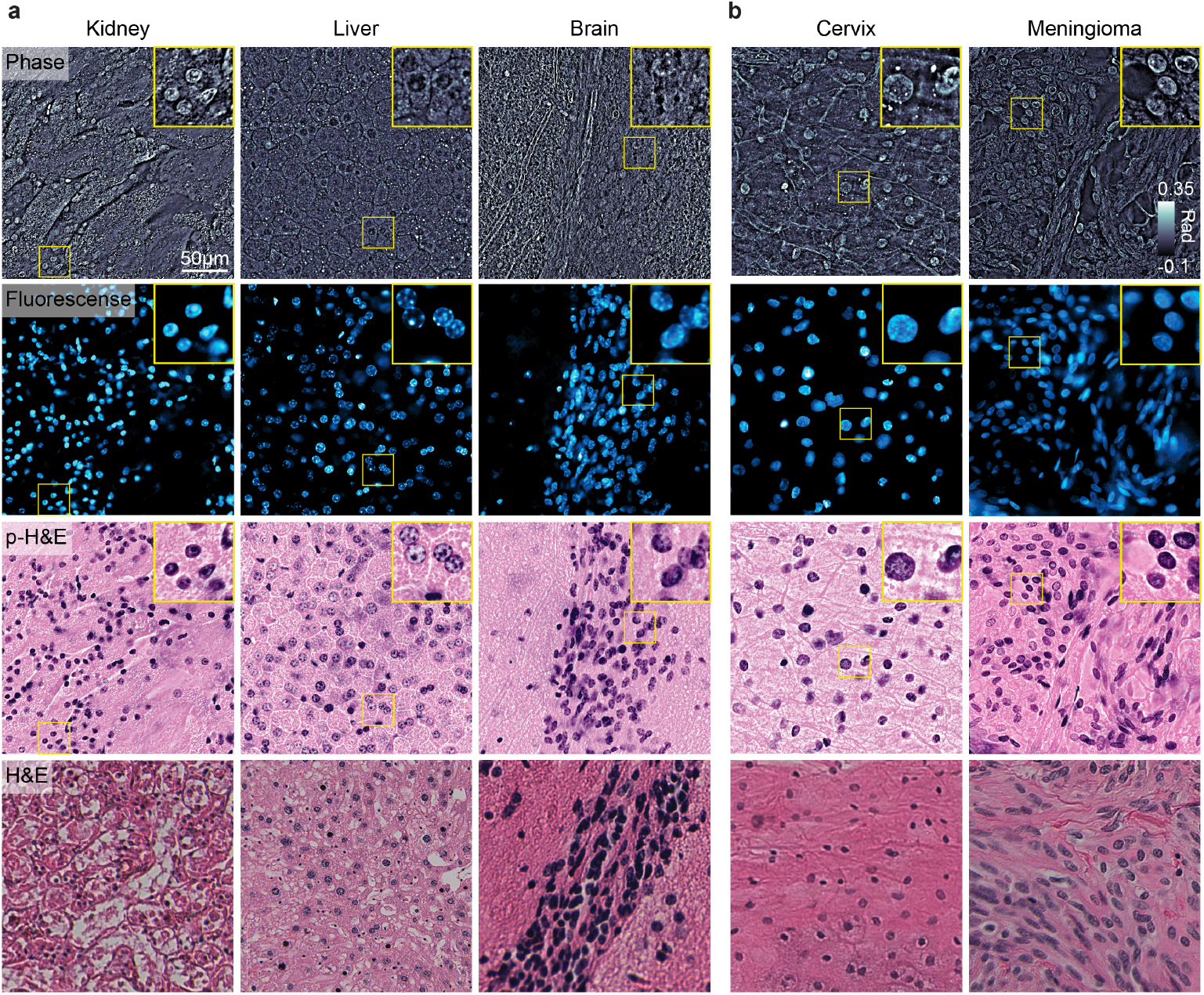
Representative multimodal and pseudo-H&E (p-H&E) images of various tissues along with real H&E stained thin tissue sections of the same tissues in similar areas. **(a)**: Different mouse organs imaged by the multimodal system. The rows are: qOBM phase images, DAPI fluorescence images, pseudo-H&E (p-H&E) images generated from the combined phase and fluorescence data, and standard histopathology (H&E) for comparison. **(b)**: Images of human meningioma and human cervix tissue.

### 2.2 Slide-free pseudo-H&E

Next, we demonstrate that the co-registered qOBM phase and DAPI fluorescence images can be used to generate pseudo-H&E images with high fidelity across different tissue types. Figure 2 presents example pseudo-H&E images from multiple organs, including both healthy and cancer specimens from mouse (2a) and human (2b). The human tissues were de-identified and discarded surgical specimens provided by the Emory University Hospital, and the mouse specimens were obtained from animals that underwent euthanasia for unrelated IACUC-approved studies at Georgia Institute of Technology. All tissues were imaged within ∼2 hours of extraction.

By using the DAPI fluorescence channel as a hematoxylin-like nuclear stain (purple) and the phase image as an eosin-like counterstain (pink), we generate pseudo-H&E images of bulk tissue that closely reproduce the appearance of conventional H&E-stained sections. These images capture the features pathologists rely on in H&E slides, including nuclear size, shape, distribution, sub-nuclear structures, and even the fibrous appearance of the stroma captured by the phase image, without the time-consuming pipeline required for standard histology, which takes ∼10 hours or more. The last row of Fig. 2 shows the real H&E images from a similar region of the same bulk tissue imaged with the multimodal system, which show nearly the same structural detail as the pseudo-H&E. The slight stylistic mismatch between the real H&E images of the corresponding tissue sections and pseudo-H&E images of unprocessed bulk tissues is the result of tissue fixation and other tissue processing steps that alter the morphology of fresh tissue. Otherwise, this simple pseudo-H&E colorization faithfully recapitulates the visual contrast and structure of H&E stained tissue sections, the gold standard in histology. (Because this work focuses on the introduction of the multimodal system and its capabilities, we leave additional determinations of diagnostic equivalence of the pseudo-H&E images for future work.)

### 2.3 Virtual fluorescence staining and virtual pseudo-H&E via Deep Learning

Previous QPI-based virtual staining methods have been trained on thin sections or cultured cells imaged in transmission [5, 25, 26]. Even when such models were applied to thicker tissue sections (up to 50 *µ*m), training to virtual histology relied on thin processed sections because paired volumetric ground truth for thick samples was unavailable [26]. Given the lack of paired data for training, researchers have explored unsupervised approaches, such as CycleGAN-based methods that learn mappings between unpaired label-free and stained image datasets [27, 28]. However, these unsupervised approaches are prone to hallucination artifacts: without paired targets during training, the cycle-consistency loss can be satisfied by mappings that do not faithfully preserve the underlying structure [29].

Our multimodal system addresses these limitations by acquiring co-registered phase and fluorescence images directly from fresh, unaltered thick tissue. These pixel-to-pixel matched images allow us to train a supervised network that converts label-free quantitative phase images into virtual fluorescence images. As a result, nuclear staining is achieved without fluorescent labels, providing an efficient and cost-effective alternative to fluorescence imaging that avoids its main limitations, including limited penetration depth, phototoxicity, photobleaching, and the very need for exogenous labels and time-consuming staining protocols. Later, we can also combine the virtual fluorescence images with phase images to generate virtual pseudo-H&E images.

The dataset used for training the supervised network consists of images acquired from fresh, thick mouse brain tissue samples using our multimodal system. A total of 1300 image pairs from the surface (*z <* 30 *µ*m where tissues are well stained) were preprocessed and selected and divided into train, validation and test set (see Training set build section in Methods for details). The training was conducted on a high-performance computing system equipped with an NVIDIA GeForce RTX 3070. All presented results are from independent animal samples and tissue regions not seen by the training algorithm.

Figure 3**a** shows the training pipeline, and 3**b** shows representative virtual staining results from the trained network on test images. The network performs consistently across regions with varying cellular density and surrounding tissue structure. The inferred virtual fluorescence images also faithfully recover sub-nuclear structures. Network performance was quantified using the peak signal-to-noise ratio (PSNR) and the structural similarity index measure (SSIM), computed on a held-out test set not used during training. The network achieves an average SSIM of 0.85 and an average PSNR of 20 dB (Fig. 3**c**), indicating that the virtually stained images faithfully reproduce the structural features of the ground-truth fluorescence images, including nuclear morphology and distribution, while preserving the overall intensity distribution.

**Fig. 3.**
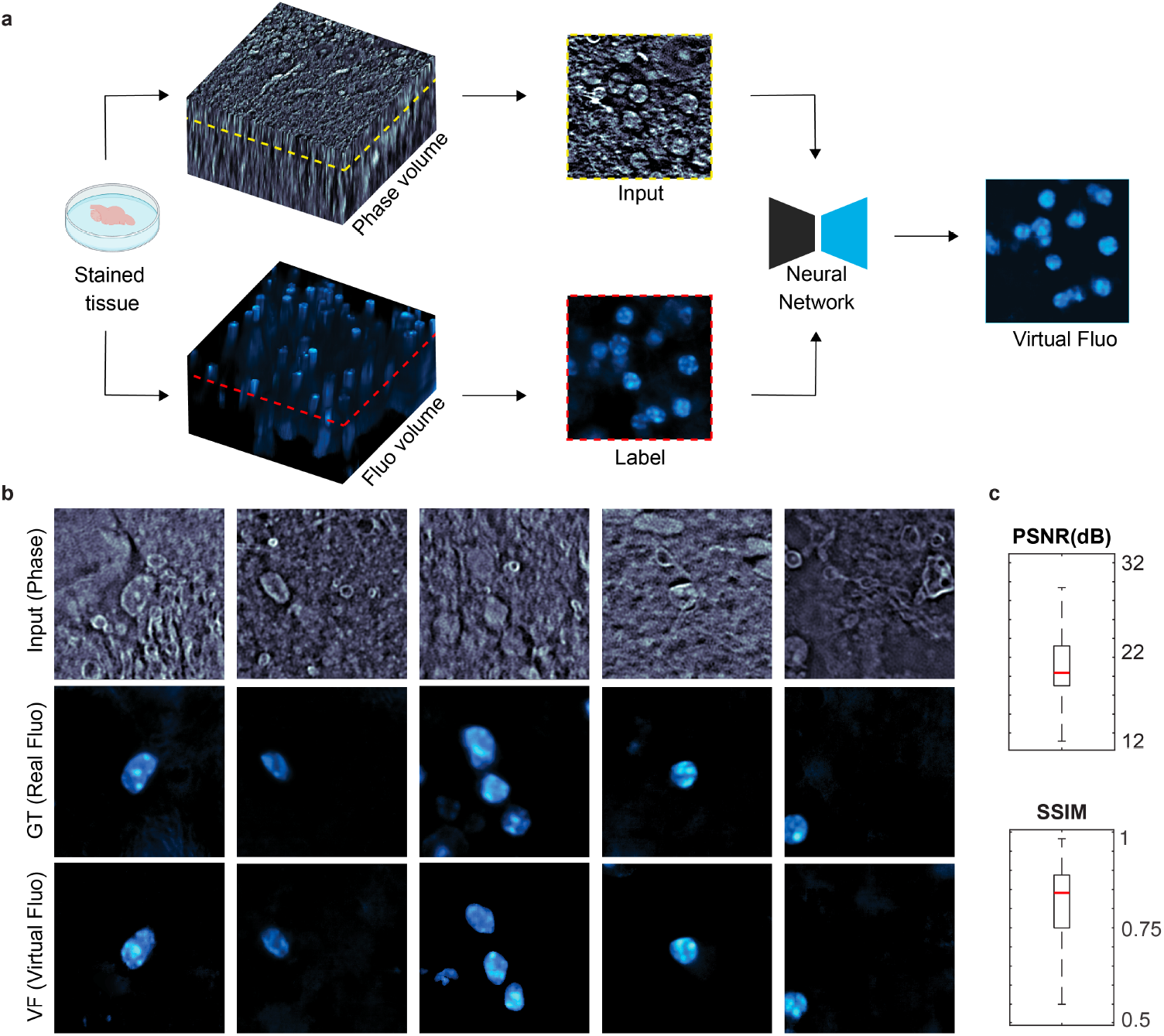
Supervised training of the virtual fluorescence network. **(a)**: Schematic of the data acquisition and network training pipeline, using co-registered qOBM phase and DAPI fluorescence images of fresh mouse brain. **(b)**: Representative qOBM phase, measured fluorescence, and virtual fluorescence images from the held-out test set. **(c)**: Peak signal-to-noise ratio (PSNR) and structural similarity index measure (SSIM) between virtual and measured fluorescence across the full test set.

As briefly mentioned, the virtual staining approach also shows important advantages over deconvolution epi-fluorescence beyond simply avoiding phototoxicity and photobleaching. Notably, although the network was trained on data acquired near the tissue surface, where the fluorescence signal is strongest and the stain is more homogeneous, the network can still generate virtual fluorescence images from the qOBM phase at depths well beyond the reach of wide-field fluorescence imaging. Figure 4**a** shows co-registered volumes of qOBM phase, real fluorescence, and virtual fluorescence, and Fig. 4**b** shows representative *x* –*y* sections of brain tissue at superficial (0 and 15 *µ*m), intermediate (50 and 75 *µ*m), and deep (130 and 140 *µ*m) planes. As imaging moves away from the surface, the measured fluorescence signal weakens and background noise increases substantially. Instead, the virtual fluorescence retains high nuclear contrast significantly deeper into the tissue.

**Fig. 4.**
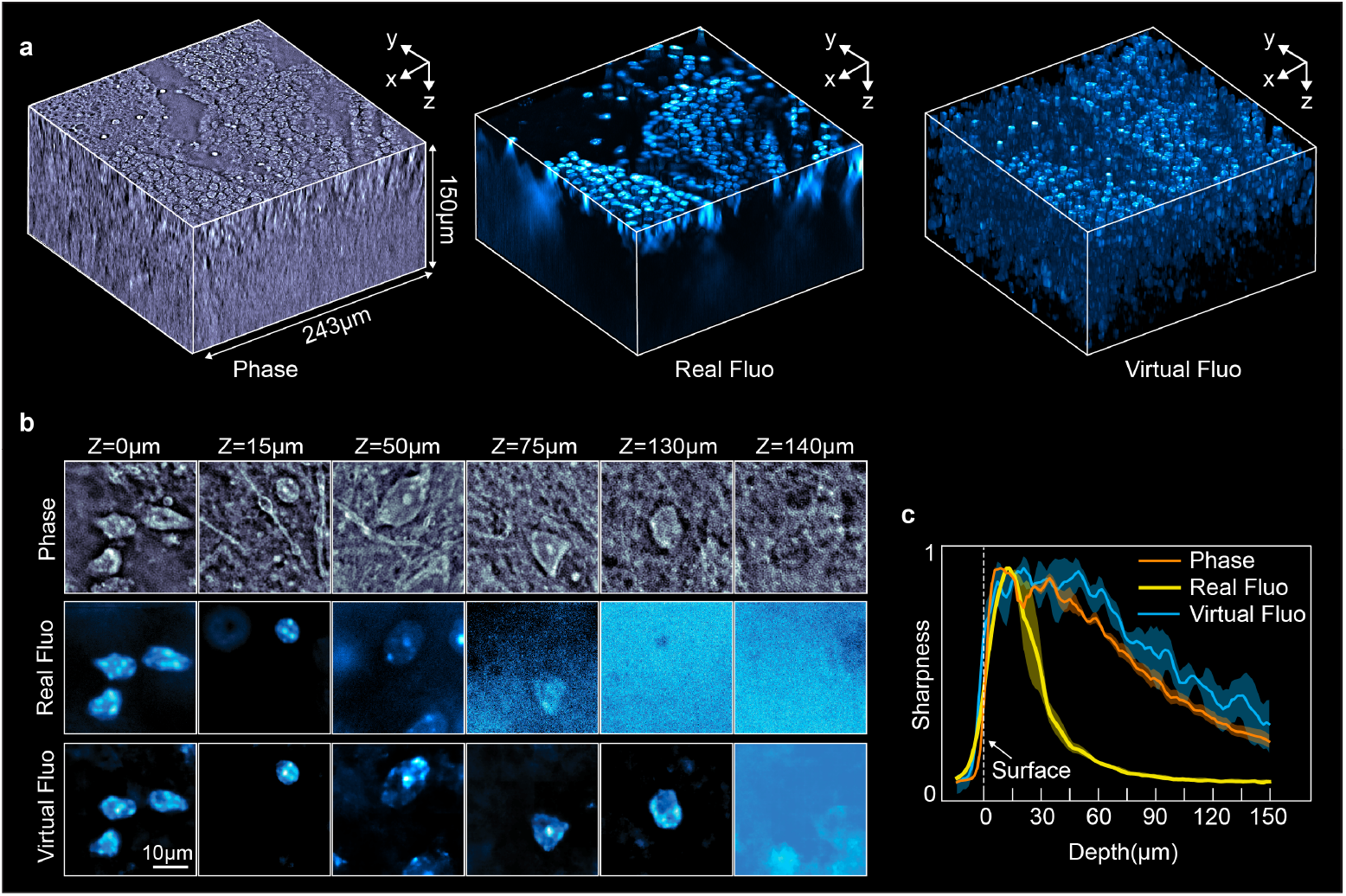
**(a)**: Three-dimensional volumes acquired with quantitative phase imaging, the corresponding DAPI fluorescence (ground truth), and the virtual staining predicted by the trained network. **(b)**: Representative *en face* images from each modality at superficial (0 and 15 *µ*m), intermediate (50 and 75 *µ*m), and deep (130 and 140 *µ*m) planes within the tissue. **(c)**: Normalized noise-corrected sharpness as a function of imaging depth for qOBM phase, measured fluorescence, and virtual fluorescence (Supplementary Section 3). Solid lines show the mean across three regions, and shaded areas indicate *±*1 standard deviation at each depth.

To quantify the imaging depth improvement, we computed a noise-corrected sharpness metric for each layer of the phase, real and virtual fluorescence volumes. The metric is the Sobel sharpness of each image, minus the contribution expected from that image’s noise level, so that a noise-only image scores approximately zero (see Supplementary Section 3 for details). The same metric was applied to all three modalities across three volumes. As shown in Fig. 4**c**, real fluorescence loses nuclear contrast rapidly with depth: it falls to half of its maximum at ∼30 *µ*m and becomes very noisy at approximately 60 *µ*m. In contrast, both the qOBM phase and the virtual fluorescence retain high contrast beyond 130 *µ*m. Because the virtual staining is inferred from the phase, its decline with depth follows that of the phase signal, indicating that its depth penetration is limited by the phase image rather than by the more limited fluorescence excitation and collection. We note that these results are representative of brain tissue–other tissue types will result in different penetration depths, depending on their absorption and scattering properties. Nonetheless, the use of longer wavelengths by qOBM will inevitably enable deeper imaging with more homogeneous virtual staining at depth than wide-field fluorescence.

Finally, we combine the qOBM phase and virtual fluorescence volumes to generate virtual pseudo-H&E (vp-H&E). The pipeline is illustrated in Fig. 5**a**: first a qOBM phase volume is acquired, then the trained network converts the phase volume into a virtual fluorescence volume, and the two volumes are finally combined into a single vp-H&E volume. Because thick, fresh tissue requires no staining and inference takes approximately one second per 2048 × 2048-pixel phase image, this pipeline substantially shortens the time required for histology and eliminates the need to physically stain the sample.

**Fig. 5.**
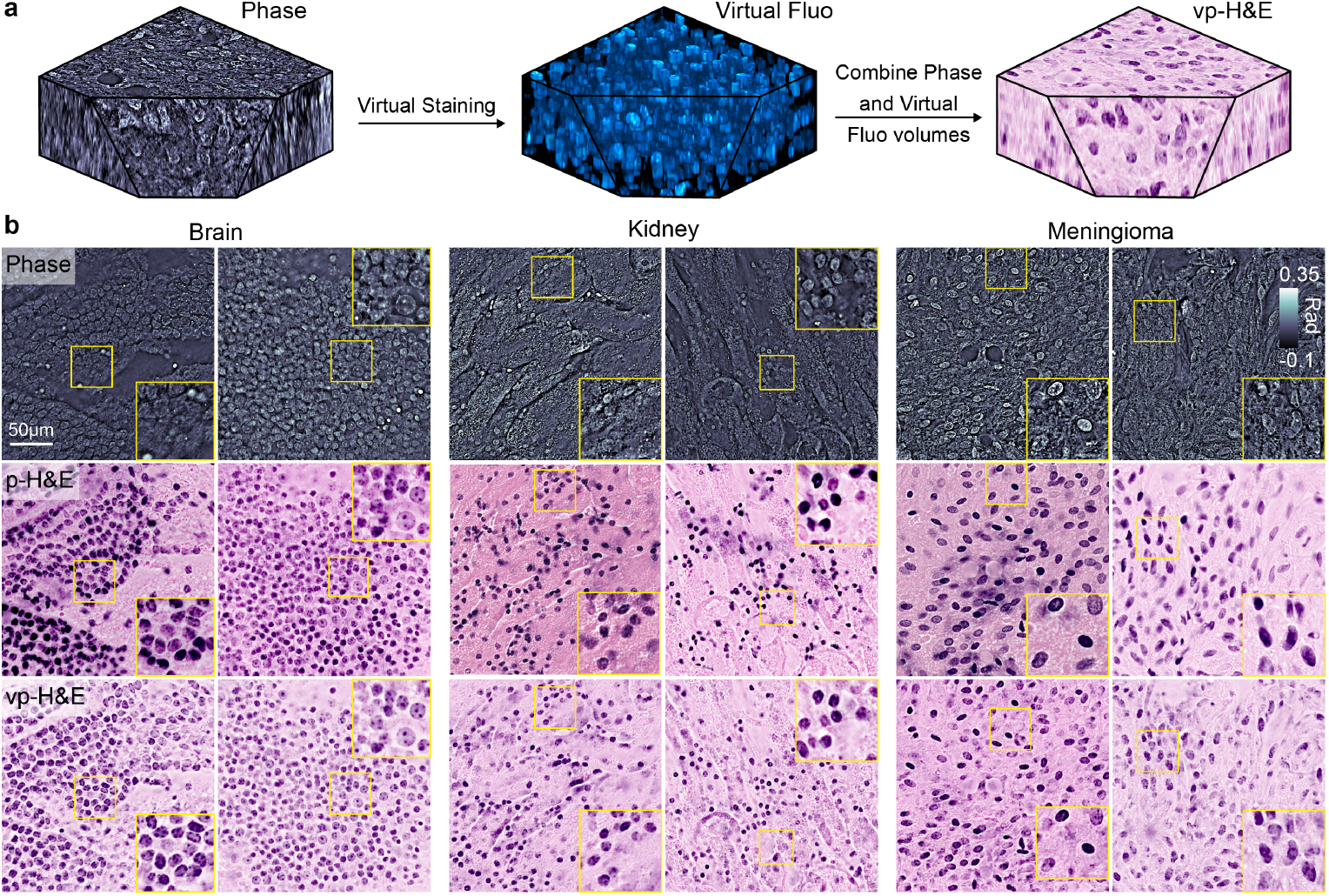
**(a)**: Schematic of the proposed pipeline, in which quantitative phase serves as the sole input modality for generating H&E-like images. **(b)**: Representative images acquired with a 60*×* objective from brain, kidney, and meningioma tissue. Top row: quantitative phase images. Middle row: pseudo-H&E images rendered by combining phase with experimentally acquired DAPI fluorescence. Bottom row: virtual pseudo-H&E images generated from phase alone using the proposed pipeline.

Figure 5**b** shows additional vp-H&E results, alongside the corresponding qOBM phase images and pseudo-H&E (p-H&E) images generated from real fluorescence. In mouse brain (left), vp-H&E agrees well with the p-H&E nuclear distribution, even in densely nucleated regions. To assess generalization, we further apply the network, without any retraining, to tissues not seen during training. In mouse kidney (middle), the network produces nuclear contrast consistent with the real staining with SSIM = 0.62 and 0.70, PSNR = 17.8 and 18.05 dB for the two examples shown. This robust performance in previously unseen tissue types is likely because it learns nuclear shape and structural features from the phase images rather than organ-specific context.

In human meningioma (right), a tumor arising from the meninges that surround the brain and spinal cord, nuclei are not only larger than those in healthy mouse brain but also vary considerably in shape, ranging from elongated to ovoid. Despite never having seen this tissue type during training, the network captures this morphological variability, faithfully reproducing both elongated and ovoid nuclei (SSIM = 0.64 and 0.56, PSNR = 20.82 and 16.63 dB). These zero-shot results indicate that features learned from healthy mouse brain transfer well to other tissue types, including human tumor specimens. For tissues whose nuclear morphology or distribution differs more substantially, such as mouse liver, direct application is less effective (see Supplementary Fig. S3). However, because the network already captures general nuclear features, we expect that it can be efficiently adapted to such cases, including clinical histology specimens, through transfer learning with only a small amount of additional paired data, as has been demonstrated for other label-free virtual staining modalities [30, 31].

## 3 Discussion

In this work, we presented a multimodal microscope that combines quantitative oblique back-illumination microscopy (qOBM) with wide-field epi-fluorescence deconvolution microscopy to acquire intrinsically co-registered volumetric phase and fluorescence images of thick, fresh tissue. We demonstrated that fluorescence and qOBM phase images can be combined to produce pseudo-H&E images of fresh bulk tissue, which provides a path towards 3D slide-free histology without the use of AI and with only one stain. We also leveraged the pixel-wise co-registered phase and fluorescence volume pairs to train a supervised deep learning neural network to predict nuclear fluorescence directly from the qOBM phase images. The qOBM phase-derived virtual fluorescence provides the advantages of both modalities, namely high nuclear contrast without exogenous stains, photobleaching, or phototoxicity, and–because qOBM uses longer wavelengths–it also enables virtual fluorescence imaging at deeper depths. The clear subcellular detail of qOBM, the simplicity of the multi-modal system, and also the ability to provide label-free virtual fluorescence are significant improvements compared to recent state-of-the-art fluorescence and RI tomography methods [23]. Finally, by combining the phase and virtual fluorescence volumes, we can also generate virtual pseudo-H&E volumes of fresh thick tissue without any staining.

This two-step design, which first predicts nuclear fluorescence using a supervised algorithm and then combines it with the phase image to generate pseudo-H&E, offers several advantages over one-step, unpaired approaches such as CycleGAN-based virtual H&E staining [28, 32]. The intermediate virtual fluorescence is trained with pixel-wise supervision and can be validated directly against measured fluorescence, while the final colorization follows a deterministic mapping rather than a learned style transfer. Together, these features enable direct comparison of the network output with the nuclear ground truth and reduce the risk of hallucinated structures, a known limitation of unpaired image translation.

It is important to note that we chose DAPI as the nuclear stain because, among the nuclear dyes we tested (including SiR-DNA and Hoechst), it caused the smallest disruption to the refractive index composition of the tissue and thus phase contrast. In other words, the tissue appeared similar before and after labeling with DAPI (see Supplemental Section 2). With Hoechst, for example, the high concentrations and long incubation times required for bulk staining produced inhomogeneous staining and noticeable non-nuclear signals. Also, the RI distribution and hence phase contrast were significantly altered after staining (Fig. S2). This is important because the trained network is intended to be applied to unstained tissue, so the phase images used for training need to closely match those of unstained samples. However, a limitation of this study lies in the quality of the DAPI ground-truth fluorescence data. In fresh, bulk tissue, DAPI does not stain evenly, since the dye penetrates incompletely and is taken up differently by different tissues, so some nuclei are missed or only weakly stained, especially deeper in the sample. This could be improved in the future by carefully optimizing the type of stain and protocols that provide a more even staining, while also causing minimal alterations to the refractive index composition of the stained tissue.

While wide-field deconvolution microscopy reduces out-of-focus background considerably, it does not match the optical sectioning capabilities of confocal or two-photon microscopy, so some blur and background signals remain present in the “ground truth”. Because of these imperfections, the training data had to be carefully curated to exclude poorly stained or low-contrast regions, and the loss function had to be designed to take into consideration miss staining.

Finally, the present study focused on single-channel nuclear fluorescence (DAPI), but the system readily supports multi-channel fluorescence acquisition, enabling virtual staining of additional cellular components and molecular markers. Looking forward, we envision extending this platform toward multiplexed molecular imaging by training models to predict the distributions of multiple organelles and molecular species from quantitative phase data alone. Recent work on label-free prediction of landmark organelles in 2D cell cultures supports the feasibility of this approach [33], but importantly, our system extends this capability to volumetric imaging of fresh, unaltered bulk tissue. Such capabilities could benefit applications such as intraoperative imaging, where rapid, label-free tissue assessment could guide surgical decision-making. By combining co-registered multimodal imaging with supervised deep learning, this platform represents a step toward label-free, high-contrast, structure-specific volumetric microscopy *in vivo* for a wide range of biomedical and clinical applications.

## 4 Methods

### 4.1 Staining and imaging protocols

Freshly excised tissue samples were stained with DAPI (4’,6-diamidino-2-phenylindole; Boster, AR1176) to label cell nuclei. Healthy tissues, such as mouse brain and liver, were stained for 1 h at a concentration of 1 *µ*g/mL. This concentration and duration were higher and longer than the protocols used for staining cells on slides, to allow the dye to diffuse deeper into the tissue, but not too long to alter the tissue morphology or RI composition. To confirm that staining did not perturb the sample, we compared the refractive index and morphology of brain tissue before and after staining and observed no noticeable changes (see Supplementary Fig. S2). For tumor tissues, such as human meningioma samples, an extended staining duration was employed (*>*2 hours). This adjustment accounted for the unique properties of cancer cells, which often have increased membrane thickness and rigidity compared to normal cells [34], which slow passive dye uptake. After staining, samples were washed three times with phosphate-buffered saline (PBS) to remove residual dye.

During qOBM imaging, the four LEDs were switched sequentially at 12 Hz with a typical exposure time of 50 ms. Fluorescence images were acquired immediately after each phase acquisition, ensuring that both modalities captured the same field-of-view (FoV) and depth. Fluorescence exposure times depended on the sample: 450 ms (2 Hz) for healthy tissue and 900 ms for tumor samples, which exhibited weaker fluorescence signal. To improve the SNR, each FoV was imaged three times and averaged. Although more averages would further improve SNR, we limited averaging to three frames to minimize photobleaching.

### 4.2 Fluorescence data processing

Raw fluorescence volumes were pre-processed in two steps, background normalization and 3D deconvolution, to provide high-fidelity ground truth. Because the samples were thick, stained tissues, fluorescence from nuclei above and below the focal plane contributed substantial background and caused uneven intensity across the FoV (Supplementary Fig. S1**b**). This effect was strongest in regions of high nuclear density, such as the pyramidal cell layer of the hippocampus. To correct it, we acquired a background image by translating the sample to an out-of-focus position where individual structures were no longer resolved, and smoothed the blurred fluorescence signal with a 50-pixel averaging window to create a normalization map. Each slice of the raw fluorescence volume was then divided by this map, yielding uniform intensity across the lateral FoV (Supplementary Fig. 1**c**).

To further suppress out-of-focus light inherent to wide-field microscopy and improve axial resolution, we deconvolved the normalized volumes using DeconvolutionLab2 [35] with the Richardson-Lucy algorithm (10 iterations). The point spread function was generated with the Born & Wolf 3D model using an objective with 0.7 NA and 461 nm emission wavelength. Deconvolution enhanced sub-nuclear detail and optical sectioning, improving structural correspondence with the qOBM phase data (Supplementary Fig. 1**d**).

### 4.3 Training set build

Training data were acquired from 12 mouse brains, each cut in half to expose a larger imaging surface. Again, we chose DAPI as the nuclear stain because it causes little to no change to tissue refractive index, morphology and stiffness, unlike other stains we tested (SiR-DNA and Hoechst). Thus, qOBM phase images of stained and unstained tissues looked the same and had similar phase distribution, which is critical for ensuring that the algorithm trained with stained images can robustly virtually stain images of unstained tissue. However, DAPI did not stain all nuclei in fresh tissue, as some nuclear membranes remained intact.

To reduce the influence of unstained nuclei, we curated the dataset manually. Paired images were divided into 256 × 256-pixel patches, and only patches containing predominantly in-focus, well-stained nuclei were retained. Patches with good phase contrast but poor staining were discarded. The curated patches were then split into training, validation, and test sets (70%, 20%, and 10%). Test set data were from organs that were completely held out from training. Data augmentation was applied to have a total of approximately 1300 images.

### 4.4 Network training

The network used a generative adversarial network (GAN) framework [36], with a U-Net [37] generator and a convolutional neural network (CNN) classifier as the discriminator. U-Net is widely used for biomedical image translation [30, 38, 39] because its skip connections preserve fine structural detail, while adversarial training encourages realistic fluorescence-like outputs.

The standard pix2pix loss [40] relies on a pixel-wise L1 term, which heavily penalizes the network whenever it predicts a nucleus that is visible in the phase image but missing from the imperfectly stained ground truth. To make training more robust to such cases, we added focal loss [41], Dice loss [42], and a structural similarity (SSIM) loss to the objective. Focal loss reweights pixels according to prediction difficulty, Dice loss emphasizes nuclear shape and regional overlap, and the SSIM loss encourages preservation of structural detail. The total loss was

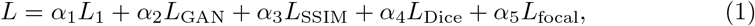

where *α*_1_–*α*_5_ are weighting coefficients.

## 5 Author contributions

ZL and FER conceived the study and designed the system. SB performed experiments, analyzed the data. ADST helped with sample preparations. DH helped with the system design. KN performed experiments and data analysis. NP helped design and validate the network. JPC helped with sample preparation and performed experiments. ZL and FER wrote the manuscript. All authors reviewed and approved the final manuscript.

