## Supplemental file for "Multimodal epi-mode 3D quantitative phase and fluorescence microscopy enables supervised volumetric virtual fluorescence in bulk tissue"

### Supervised 3D Virtual Fluorescence and Pseudo-H&E Staining of Fresh Bulk Tissues via Multimodal qOBM and Fluorescence Microscopy

Zhenmin Li[1*], Srinidhi Bharadwaj[2], Aaron D Silva Trenkle[2], Daniel Houston[2], Kyan Nguyen[2], Nimai Ponna[1], Jacob P. Callaway[2], Gabriel A. Kwong[2], Francisco E. Robles[2,3*]

[1]Electrical and Computer Engineering, Georgia Institute of Technology, Atlanta, 30332, GA, USA.
[2]Wallace H. Coulter Department of Biomedical Engineering, Georgia Institute of Technology and Emory University, Atlanta, 30332, GA, USA.
[3]Department of Imaging Sciences, St. Jude Children's Research Hospital, Memphis, 38105, TN, USA.

### Supplementary information

#### 1 Fluorescence pre-processing and deconvolution

This section describes how raw fluorescence images acquired with the multimodal qOBM system were processed before training. When imaging densely nucleated regions of thick tissue, such as the mouse hippocampus, out-of-focus fluorescence produces uneven background across the field of view (see Fig. S1**b**). To correct this, we first acquired an image of the same field of view with the sample translated out of focus and then smoothed it with a $50 \times 50$-pixel averaging window to obtain a background map (Fig. S1**a**). Next, each raw image was divided by this map to homogenize intensity across the field of view (Fig. S1**c**), and then the background normalized volume was deconvolved in 3D using Richardson–Lucy deconvolution algorithm to improve the contrast (Fig. S1**d**). The 3D point spread function (PSF) used was Born & Wolf 3D model with a wavelength at 450nm.

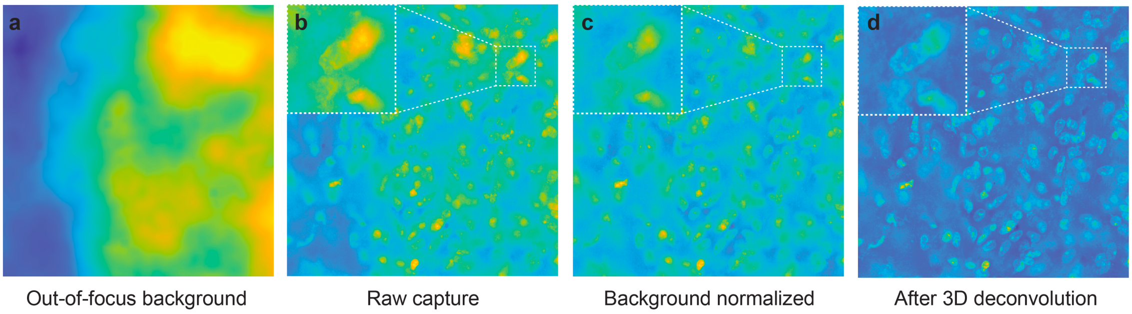


**Fig. S1** | Fluorescence pre-processing pipeline. **(a)**: Normalization map obtained from an out-of-focus image smoothed with a $50 \times 50$-pixel window. **(b)**: Raw fluorescence image. **(c)**: Fluorescence image after normalization. **(d)**: Final fluorescence image after 3D Richardson–Lucy deconvolution.

#### 2 Comparison of nucleic acid dyes

In this section, we compare the effects of different nuclear stains on fresh mouse brain tissue (Fig. S2). We tested SiR-DNA (3 $\mu$M, 2 h), Hoechst 33342 (10 $\mu$g/mL, 30 min), and DAPI (1 $\mu$g/mL, 1 h), with concentrations and incubation times chosen to yield comparable fluorescence signal and contrast. Among the three dyes, DAPI produced the smallest change in phase contrast relative to fresh, unstained mouse brain and showed no detectable signal outside nuclei. In contrast, SiR-DNA and Hoechst produced larger changes in phase contrast and stained non-nuclear structures, and Hoechst still left some nuclei unstained. We therefore selected DAPI as the nuclear stain for this work.

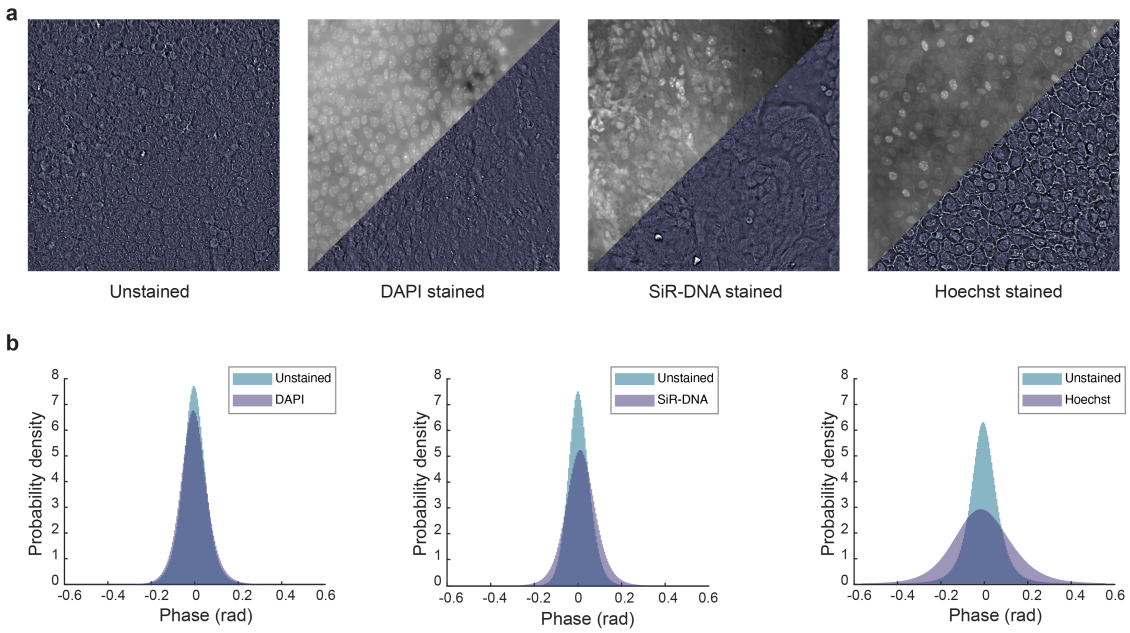


**Fig. S2** | **(a)**: qOBM images and fluorescence images of unstained, DAPI stained, SiR-DNA stained and Hoechst stained mouse brain. **(b)**: Probability density of phase values of fresh unstained mouse brain plotted with the probability density of phase values after DAPI, SiR-DNA and Hoechst staining of the same mouse brain.

### 3 Depth-resolved sharpness metric

To compare how well nuclear structure is preserved with depth across the qOBM phase, measured fluorescence, and virtual fluorescence volumes, we used a noise-corrected sharpness metric, applied identically to all three modalities. For each image, we first computed the raw sharpness $S$, which is the Sobel gradient of the image. Because noise also contributes to $S$ and boosts the value for real fluorescence images when getting deeper into tissue, we estimated the noise level $\sigma_n$ using Immerkær's method [1] and subtracted its expected contribution to flatten the curve for the real fluorescence.

$$\tilde{S} = S - c\,\sigma_n, \tag{1}$$

where $c = 0.057$ is a constant, applied so a layer containing only noise scores approximately zero.

### 4 Direct inference on mouse liver

Here we show the results of applying the network, trained only on mouse brain, directly to fresh mouse liver without retraining (Supplementary Fig. S3). As noted in the main text, direct application to liver performed poorly. Fig. S3**a** shows a region near the surface of the intact liver, and Fig. S3**b** shows a deeper region with the morphology typical of liver parenchyma. At the surface, some individual liver nuclei slightly resemble the phase appearance of those in mouse brain: they appear brighter than their surroundings in the phase image. Here, inference is slightly better (SSIM = 0.43), and several nuclei are correctly converted (yellow and orange arrowheads). In the deeper region, however, nuclei appear darker in the cells in the phase image, surrounded by bright, high RI lipids. This contrast is the reverse of what the network learned from brain tissue, and the network fails to identify nuclei (SSIM = 0.33).

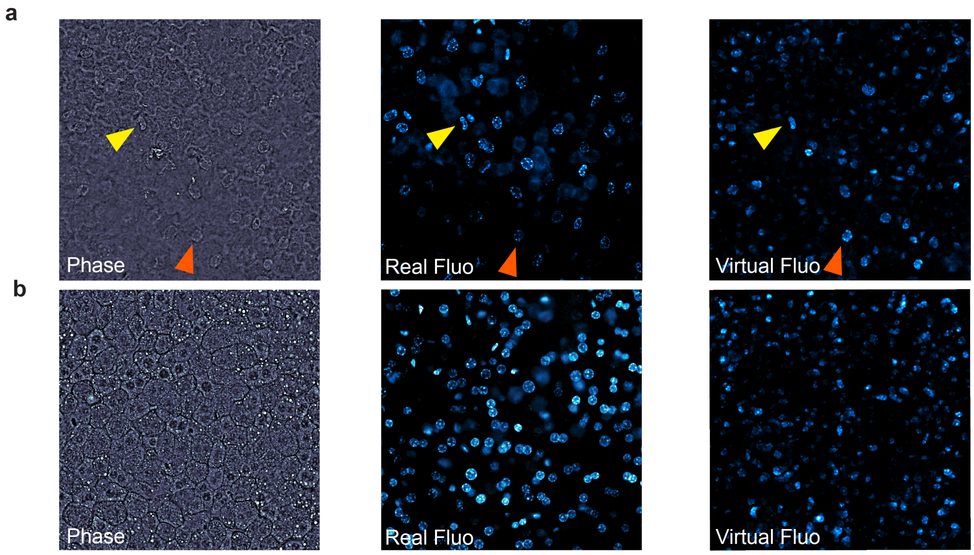


**Fig. S3** | **a**: Region near the liver surface, where nuclei appear brighter than their surroundings in the qOBM phase image; arrowheads mark nuclei correctly reproduced by the network. **b**: Deeper region with typical liver morphology, where nuclei appear darker. Columns show qOBM phase, measured fluorescence, and virtual fluorescence.
